# Nitrogen Fertilisers Shape the Composition and Predicted Functions of the Microbiota of Field-Grown Tomato Plants

**DOI:** 10.1101/672162

**Authors:** Federica Caradonia, Domenico Ronga, Marcello Catellani, Cleber Vinícius Giaretta Azevedo, Rodrigo Alegria Terrazas, Senga Robertson-Albertyn, Enrico Francia, Davide Bulgarelli

**Affiliations:** Department of Life Sciences, Centre BIOGEST-SITEIA, University of Modena and Reggio Emilia, Italy; Faculty of Agricultural and Veterinary Sciences, Sào Paulo State University, Jaboticabal, Brasil; Plant Sciences, School of Life Sciences, University of Dundee, Dundee, United Kingdom

**Author notes:** CREA, Centre for Animal Production and Aquaculture (CREA-ZA), Lodi, Italy. ENEA, Department for Sustainability, C.R. Trisaia, Rotondella (MT), Italy. Correspondence: Federica Caradonia, Enrico Francia, Davide Bulgarelli.

**Keywords:** *Solanum lycopersicum*, Rhizosphere, Root, Microbiota, Nitrogen, Fertilisers, Digestate

## Abstract

The microbial communities thriving at the root-soil interface have the potential to improve plant growth and sustainable crop production. Yet, how agricultural practices, such as the application of either mineral or organic nitrogen fertilisers, impact on the composition and functions of these communities remains to be fully elucidated. By deploying a two-pronged 16S rRNA gene sequencing and predictive metagenomics approach we demonstrated that the bacterial microbiota of field-grown tomato (*Solanum lycopersicum*) plants is the product of a selective process that progressively differentiates between rhizosphere and root microhabitats. This process initiates as early as plants are in a nursery stage and it is then more marked at late developmental stages, in particular at harvest. This selection acts on both the bacterial relative abundances and phylogenetic assignments, with a bias for the enrichment of members of the phylum Actinobacteria in the root compartment. Digestate-based and mineral-based nitrogen fertilisers trigger a distinct bacterial enrichment in both rhizosphere and root microhabitats. This compositional diversification mirrors a predicted functional diversification of the root-inhabiting communities, manifested predominantly by the differential enrichment of genes associated to ABC transporters and the two-component system. Together, our data suggest that the microbiota thriving at the tomato root-soil interface is modulated by and in responses to the type of nitrogen fertiliser applied to the field.

## INTRODUCTION

Limiting the negative impact of agricultural practices on the environment while preserving sustainable crop yield is one of the key challenges facing agriculture in the years to come.

As an essential element for plant nutrition, nitrogen represents a paradigmatic example of such a challenge. Moreover, due to the combined effect of elevated solubility and little retention in soils, the lack of this element is and will be one of the major yield-limiting factors worldwide (Tilman et al., 2011). At the same time, the application of synthetic nitrogen fertilisers is, in many agricultural systems, a low efficiency approach which has been linked with the degradation of natural resources (Elser and Bennett, 2011). One of the strategies adopted to limit the economic and environmental footprint of crop production while maintaining sustainable yield is the “recycling” of, mineral-rich, biodegradable products of the livestock and agricultural sectors.

One example of this approach is the application of the digestate, a by-product of the anaerobic digestion of organic waste for the production of biogas (Möller and Müller, 2012) as renewable soil amendment for crop production. The digestate is a mixture of partially-degraded organic matter, microbial biomass and inorganic compounds (Alburquerque et al., 2012). We recently demonstrated how the digestate can be efficiently used as innovative fertiliser and plant growing media (Ronga et al., 2018b; Ronga et al., 2018a; Ronga et al., 2019), yet the impact of digestate applications on the agroecosystem remains to be fully elucidated.

For instance, the digestate can be a source of phytoavailable nitrogen, in particular ammonium, capable of impacting on organic matter mineralisation and emission of carbon dioxide from the soil profile (Grigatti et al., 2011). Therefore, it is legitimate to hypothesize that such treatments impact on the composition of the microbial communities thriving at the root-soil interface, collectively referred to as the rhizosphere and root microbiota, which play a critical role in mobilisation of mineral elements for plant uptake (Alegria Terrazas et al., 2016). Congruently, several studies indicate that the application of biogas by-product enhances soil microbial activity (Möller, 2015) and the availability of phytohormones (Scaglia et al., 2015). However, the intertwined relationship among microbiota composition, soil characteristics and amendments as well as host plant species-specificity (Bulgarelli et al., 2013) makes it difficult to infer first principles.

This is particularly true for field-grown crops such as tomato (*Solanum lycopersicum* L.), one of the most cultivated horticultural crops globally with plantations occupying an area of 4.8 million of hectares with a production of 182 million tonnes in 2017 (FAO 2017). Notably, this species is also an excellent experimental model for basic science investigations: tomato was one of the first crops whose genome was sequenced (Consortium, 2012) and provided a superb platform to test the significance of genome editing for evolutionary studies and plant breeding (Zsögön et al., 2018). Perhaps not surprisingly, tomato is gaining momentum as an experimental system to study host-microbiota interactions in crop plants. Recent investigations revealed novel insights into the assembly cues of the microbiota associated to this plant (Bergna et al., 2018; Toju et al., 2019) and the contribution of microbes thriving at the tomato root-soil interface to pathogen protection (Chialva et al., 2018; Kwak et al., 2018). However, the composition and functional potential of the tomato microbiota and their interdependency from nitrogen fertilisers remain to be elucidated.

Here we report the metagenomics characterisation of the microbiota thriving at the root-soil interface of field-grown tomato plants. We hypothesize that nitrogen treatments shape and modulate the contribution of the tomato microbiota for crop yield. To test this hypothesis, we focused on processing tomato exposed to different nitrogen fertilisers, either digestate-based or containing a mineral fraction. By using a 16S rRNA amplicon sequencing survey we deciphered how the microhabitat (i.e., either rhizosphere or root) sculpts the tomato microbiota which, in turn, is fine-tuned by the type of fertiliser applied. Finally, by using a predictive metagenomics survey, we inferred the functional diversification imposed by the nature of the fertilisers on the root microbiota.

## MATERIALS AND METHODS

### Field site

A field trial was established in a tomato farm near the city of Ravenna (44°25’40.8”N 12°05’53.3”E), Emilia Romagna Region, Italy, during the 2017 growing season. During the period from transplant to harvest, the minimum and the maximum average temperatures recorded were 17.1°C and 32.8°C, respectively, and the rainfall was 101.7 mm. The soil had a silty loam texture (14% clay, 51% silt, 35% sand), a pH 8.3 (in H_2_O), 1.1 g kg^−1^ total N (Kjeldahl method), 7 mg kg^−1^ available P (Olsen method), 129 mg kg^−1^ exchangeable K (Ammonium acetate), and 9 g kg^−1^ organic matter (Walkey-Black method). A schematic illustration of the field trial is depicted in Supplementary Figure S1.

### Plant material

We used the tomato cultivar ‘Fokker’, a processing-type genotype with blocky fruit, late fruit ripening and suitable for tomato puree, for the experimentation. Seedlings were provided by Bronte Soc. Coop. Agr. A.R.L. (Mira, Italy). Processing tomato seedlings were transplanted at the end of May when they were 6-week old corresponding to plants at the fourth true leaf stage. Plant density was 3 plants m^−2^. Plants were transplanted into single row, with a spacing of 0.22 m between plants in each row and 1.50 m between rows.

### Experimental design

We established a randomized complete design with three replicates and seven treatments: pelleted digestate (hereafter PE), liquid digestate (LD), slow-acting liquid digestate (SRLD), organo-mineral fertiliser based on digestate (SC), synthetic fertiliser (MF), slow-acting synthetic fertiliser (SRMF), and no fertilization treatment (NT). The composition of the treatments is summarised in Table 1. For each treatment, we applied a total amount of nitrogen in the ratio 150 N kg ha^−1^ on the basis of soil analysis, crop rotation and crop nutrients required. Nitrogen was supplied at transplanting time with the exception of the synthetic fertiliser treatment. For this latter treatment, the amount of total Nitrogen was equally divided and applied in 3 times (transplanting, full flowering and fruit ripening) using ammonium nitrate in the first treatment and calcium nitrate in the second and in the third ones. During the trial, 600 m^3^ ha^−1^ of irrigation water was distributed by drip irrigation to each treatment. The other soil and crop management practices were performed according to the production rules of Emilia Romagna Region, Italy. Briefly, weeds control was performed with a single treatment (on 11th June) using products based on metribuzin and propaquizafop. Sulphur and Copper were used to control phytopatogenic fungi while imidacloprid, abamectin and spinosad were used as insecticide.

**Table 1.** Composition and information on fertilisers used in this study. TOC = Total organic carbon; N = Nitrogen; P = Phosphorus; K = Potassium; H2O = water content

| Treatment | TOC% | (N)% | (P)% | (K)% | H<sub>2</sub>O% | Additional information |
| --- | --- | --- | --- | --- | --- | --- |
| **Synthetic fertiliser (MF)** |  | 41 |  |  |  | Ammonium nitrate (N 26%) and calcium nitrate (N 15%) |
| **Pelleted digestate (PE)** | 39.70 | 1.50 | 2.50 | 2.00 | 7.80 | (Pulvirenti et al., 2015) |
| **Slow acting liquid digestate (SRLD)** | 3.74 | 0.34 |  | 0.95 |  | Liquid digestate plus the nitrogen stabilizer Vizura® (BASF, 2 L ha<sup>-1</sup>), |
| **Liquid digestate (LD)** | 3.74 | 0.34 |  | 0.95 |  | EC 1.07 dS m<sup>-1</sup> and pH 8.3 |
| **Organo-mineral fertiliser (SC)** | 10.50 | 10.00 | 5.00 | 15.00 | 7.00 | Produced by SCAM Spa (Modena, Italy), based on solid digestate for the organic fraction |
| **Slow acting synthetic fertiliser (SRMF)** |  | 15.00 | 15.40 | 15.00 |  | NPK Original Gold® (Compo Expert) |

### Yield traits

At harvest we determined the marketable yield (t ha^−1^), as a weight of fully ripe fruits, and the solid soluble content (°Brix t ha^−1^) as a proxy for fruit quality. The °Brix parameter was determined using the digital refractometer HI 96814 (Hanna, Italy), while the °Brix t ha^−1^ was calculated by multiplying the hectare marketable yield by °Brix and dividing the result by 100.

### Root, Rhizosphere and Bulk soil Sampling and DNA Extraction

At transplanting time (May 2017), 5 root specimens per treatment were collected. Upon uprooting, soil particles loosely bound to roots were dislodged by hand shaking and root segments of ~ 6 cm were placed in sterile 50 mL tubes. The samples were stored in a portable cooler (~ 4°C), transported to the laboratory and immediately processed. Root specimens were incubated in 30 mL of PBS (Phosphate buffered saline) and placed on a shaker for 20 minutes in order to separate the soil tightly adhering to plant material, which we operationally defined as “rhizosphere”, from the roots. The first tubes were centrifuged for 20 minutes at 4,000 x g and the rhizosphere pelleted was collected in liquid nitrogen and stored at −80°C. The roots were moved to a new sterile tube containing 30 mL PBS and sonicated by Ultrasonics Sonomatic Cleaner (Langford Ultrasonics, Birmingham, UK) for 10 minutes (intervals of 30 seconds pulse and 30 seconds pause) at 150 W, as previously reported (Schlaeppi et al., 2014) to enrich for endophytic microorganisms. Roots were then washed in the same new buffer and dried on sterile filter paper. After few minutes, the roots were moved to 50 mL tubes and frozen in liquid nitrogen for storage at −80°C. Three independent soil samples were harvested from unplanted soil in different points of the field, frozen in liquid nitrogen and stored at −80°C. At harvest time (September 2017) the whole plants were harvested, 5 roots per treatment and 3 bulk soil samples were collected, prepared and stored like the previous samples. Frozen root samples were pulverized in a sterile mortar using liquid nitrogen prior DNA preparation. DNA was extracted from all the specimens (i.e., bulk soil, rhizosphere and pulverized roots) using the FastDNA^®^ SPIN Kit for Soil (MP Biomedicals, Solon, USA) following the instruction manual provided by manufacturer. DNA samples were diluted using 50 μL DES water and quantified using the Nanodrop 1000 Spectrophotometer (Thermo Scientific, Wilmington, United States).

#### 16S rRNA Gene Sequencing

The sequencing library was generated using primers specific (515F 5’-GTGCCAGCMGCCGCGGTAA-3’ and 806R 5’-GGACTACHVGGGTWTCTAAT-3’) for hypervariable V4 region of the 16S rRNA gene. The reverse primers included a 12-mer unique “barcode” sequences (Caporaso et al., 2012) to facilitate the multiplexing of the samples into a unique sequencing run. Individual PCR reactions were performed as previously reported (Robertson-Albertyn et al., 2017), with the exception of the concentration of the Bovine Serum Albumin, added at 10 μg/reaction, and the addition of a Peptide Nucleic Acid (PNA) blocker (PNA Bio, Newbury Park, United States) at a concentration of 0.5 μM/reaction to inhibit plastidial amplification. For each barcoded primers, three technical replicates and a no-template control (NTC) were organised and processed starting from a unique master mix. Five microliters of amplified samples and cognate NTCs were inspected on a 1% (w/v) agarose gel. Two independent sets of triplicated amplicons, displaying the expected amplicon size and lacking detectable contaminations, were combined in a barcode-wise manner and purified using the Agencourt AMPure XP kit (Beckman Coulter, Brea, United States) with a ratio of 0.7 mL AMPure XP beads per 1 mL of sample. Purified DNA samples were quantified using Picogreen (Thermo Fisher, United Kingdom) and combined in an equimolar ratio into an amplicon pool. This latter material was used for the preparation of a MiSeq run at the Genome Technology facilities of the James Hutton Institute (Invergowrie, UK) as previously reported (Robertson-Albertyn et al., 2017).

#### OTU Table Generation and pre-processing

We used QIIME, version 1.9.0 (Caporaso et al., 2010) to process the sequencing output of the MiSeq machine. Briefly, the command join_paired_ends.py was used to decompress and merged (minimum overlap 5bp) forward and reverse read FASTQ files. Next, we removed *in silico* low-quality sequencing reads and sequencing reads without the barcode information. Then, the reads were assigned to individual samples. In these analyses, the command split_libraries_fastq.py was used imposing a minimum PHRED score of 20. The resulted high-quality reads were assembled into an Operational Taxonomic Unit (OTU) table at 97% sequence identity. We used a ‘closed reference’ approach for OTU-picking using the command pick_closed_reference_otus.py. We imposed the Green Genes database version 13_5 (DeSantis et al., 2006) as a reference database to identify microbial OTUs and prune for chimeric sequences. We used SortMeRNA algorithm for OTU -picking and taxonomy assignment. Finally, OTUs whose representative sequences were classified as either chloroplast or mitochondria, as well as OTUs accruing only one sequencing read over the entire dataset (i.e., singletons), were depleted *in silico* using the function filter_otus_from_otu_table.py.

#### Data visualisation and statistical analyses

Agronomic traits were analysed by Analysis of Variance (ANOVA) using GenStat 17th (VSN International, Hemel Hempstead, UK). Means were compared using Bonferroni’s test at the 5% level.

The OTU table produced in QIIME was analysed in R using a custom script developed from Phyloseq package (McMurdie and Holmes, 2013).

Initially, the data were filtered removing the samples with less than 1,000 reads and the OTUs with less than 10 reads in at least 5% of the samples. For alpha-diversity calculation, sequencing reads were rarefied at an even sequencing deep of 18,467 reads per sample retaining 2,439 unique OTUs. The number of Observed OTUs and Chao1 index were used as richness estimators, while the Shannon index was used for evaluating the evenness. Upon inspecting distribution of the data using a Shapiro-Wilk test, the means of rhizosphere and root samples at harvest time were compared using a non-parametric Wilcoxon rank sum test. Next, we performed a non-parametric Kruskal–Wallis test independently on rhizosphere and root samples to identify significant effect of the individual treatments on the ecological indices.

For beta-diversity calculation, the original counts (i.e., not rarefied) were transformed to relative abundances and we imposed an abundance threshold to target PCR-reproducible OTUs. The differences among microbial communities of the samples were computed using Bray-Curtis index and weighted Unifrac index, with this latter index including phylogenetic information in the analysis (Lozupone and Knight, 2005). A Principal Coordinates Analysis (PCoA) was generated to visualize similarities and dissimilarities of microhabitats and treatments. In order to assess the effects of microhabitats and the treatments on the bacterial community composition, a Permutational Multivariate Analysis Of Variance (PERMANOVA) on distance matrices was implemented using the function Adonis in p a two-pronged approach. First, we assessed the effect of nursery/harvest stage on microhabitat composition. Next, we used the same test to assess the impact of the treatment on rhizosphere and root specimens at harvest stage. In the two approaches, the computed R^2^ therefore reflects the proportion of variance explained by the given factor in the group of samples tested.

Finally, original counts were used to perform a differential analysis to identify individual bacteria differentially enriched in the tested samples using DESeq2 (Love et al., 2014).

The phylogenetic tree was constructed using the representative sequences of the OTUs significantly enriched in rhizosphere and root specimens and annotated with iTOL (Letunic and Bork, 2006).

#### Functional predictions

Tax4Fun (Asshauer et al., 2015) package in R was used as a predictive tool to obtain a functional profile based on 16S rRNA gene data. Metabolic capabilities are calculated by linking the amplicon data phylogenetic and abundance profile to a set of pre-computed metabolic reference profiles, based on the KEGG Ortholog (KO) database (Kanehisa et al., 2008). The input for this analysis was an OTU table obtained with the representative sequences of the OTU table previously generated (see above), reclassified using SILVA_115 taxonomic database (Quast et al., 2013). Similar to a previously reported operational protocol (Kavamura et al., 2018), we focused our analysis in prokaryotic functional categories related to amino acid metabolism, carbohydrate metabolism, cell motility, energy metabolism, membrane transport, metabolism of terpenoids and polyketides and two-component system, trimming the rest of predicted functions from the Tax4fun output. A statistical comparison between two groups using a Welch’s t-test (Bluman, 2009) filtered at a p-value < 0.01 with Storey’s correction for false discovery rate (Storey and Tibshirani, 2003) was performed in STAMP, Statistical Analysis of Metagenomic Profiles (Parks et al., 2014).

#### Data and scripts availability

The 16S rRNA gene sequences presented in this study are available at the European Nucleotide archive under the study accession number PRJEB32219. The scripts to reproduce the statistical analysis and figures are available at https://github.com/BulgarelliD-Lab/Tomato_nitrogen. Data frames required for scripts reproducibility are included in Supplementary Database 1.

## RESULTS

### Fertiliser treatment impacts on yield and quality of processing tomato

At harvest time the two most important parameters such as marketable yield and fruit quality were measured to evaluate the effect of 7 different fertiliser performances on processing tomato (Figure 1). The fertiliser treatments had a significant effect on fresh biomass of fruits (ANOVA, Bonferroni’s test, P < 0.001). Pelleted digestate registered the best performance followed by synthetic fertiliser and slow acting liquid digestate. In addition, the different fertilisers influenced significantly also the quality of processing tomato (Figure 1) (ANOVA, Bonferroni’s test, P < 0.001).

**Figure 1.**
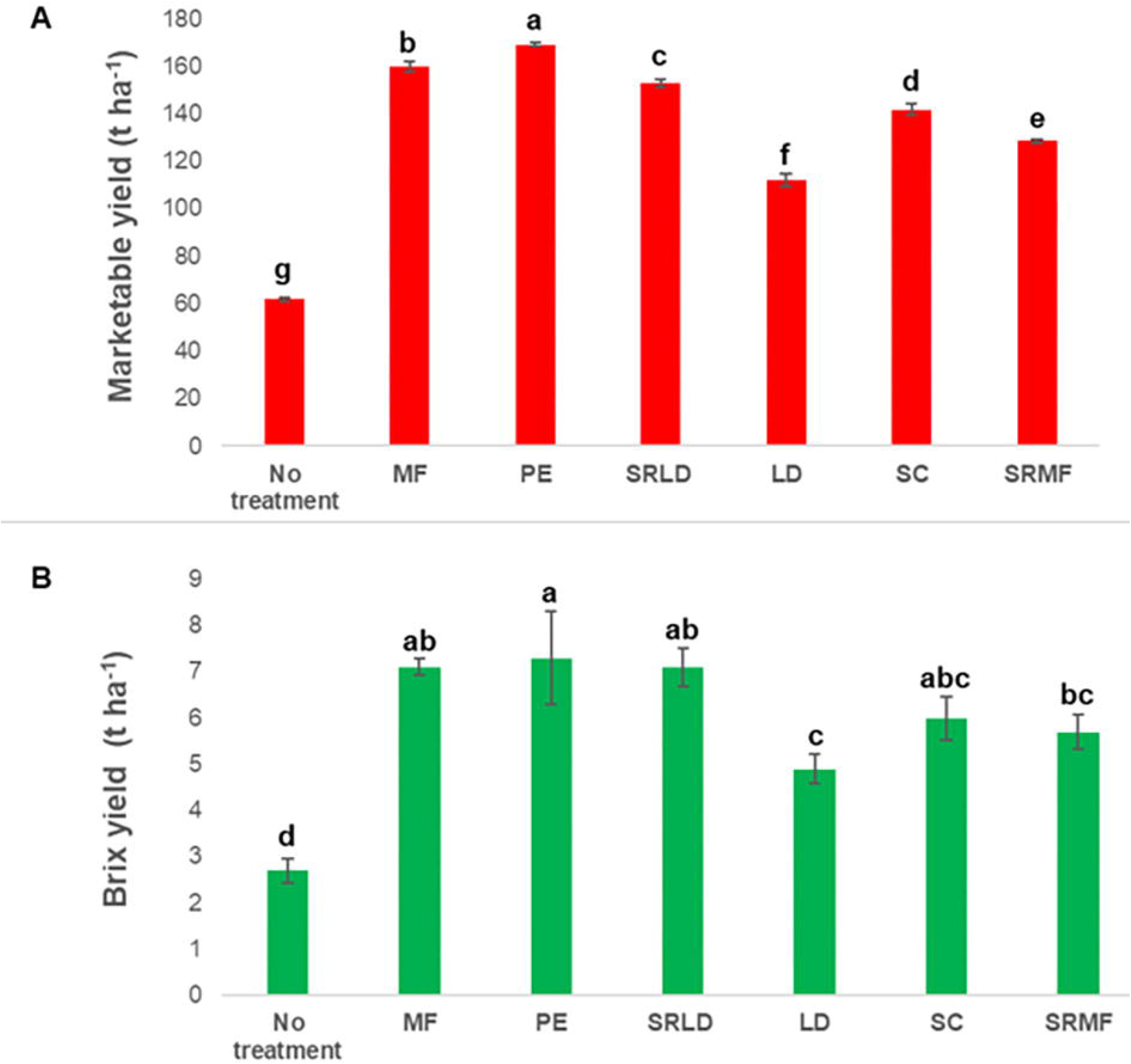
Effect of the nitrogen treatments on tomato yield traits. Mean and standard deviation of (**A**) marketable yield and (**B**) Brix of tomato plants exposed to the following treatments: LD (Liquid Digestate), SRLD (Slow acting Liquid Digestate), PE (Pelleted Digestate); SC (Organo-mineral fertiliser); MF (Mineral Fertiliser); SRMF (Slow acting Mineral Fertiliser). Different letters denote statistically significant differences between treatments by Analysis of Variance (ANOVA). Means were compared using Bonferroni’s test at the 5% level (P <0.001).

### The assembly dynamics of the bacterial microbiota of field-grown processing tomato

To gain insights into the relationships between yield traits and microbiota composition in field-grown processing tomato plants, we generated 5,546,303 high quality 16S rRNA gene sequences for the 86 samples generated in this study.

Upon *in silico* depletion of OTUs classified as Mitochondria and Chloroplast we reduced the number of analysable sequences to 4,645,503 with a retaining proportion of 83.7% of the original sequences (mean per samples = 54,017.48 reads; max = 111,213 reads; min = 272 reads). The data were further filtered removing the samples with less than 1,000 reads as well as the OTUs with less than 10 reads in 5% of samples. This allowed us to retain 2,515 unique OTUs accounting for 4,308,580 high quality reads and 85 samples.

Then, we computed alpha-diversity calculations on a dataset rarefied at 18,467 reads per sample and alpha-diversity was investigated considering two microhabitats (root and rhizosphere) and the seven fertiliser treatments. OTUs richness was assessed by Chao1 index and Observed OTUs while the OTUs evenness was assessed by Shannon index. This analysis revealed a significant effect of the microhabitat on the characteristics of the microbiota thriving at the tomato root-soil interface: regardless of the treatment, the root microhabitat emerged as less diverse and even compared to the rhizosphere one (Wilcoxon rank sum, p <0.01, Figure 2). This observation suggests that root microhabitat represents a gated community compared to the surrounding soil environment. Conversely, the treatment impacted only the number of OTUs observed in the rhizosphere compartment (Kruskal-Wallis non parametric analysis of variance followed by Dunn’s post-hoc test p < 0.05. Figure 2).

**Figure 2.**
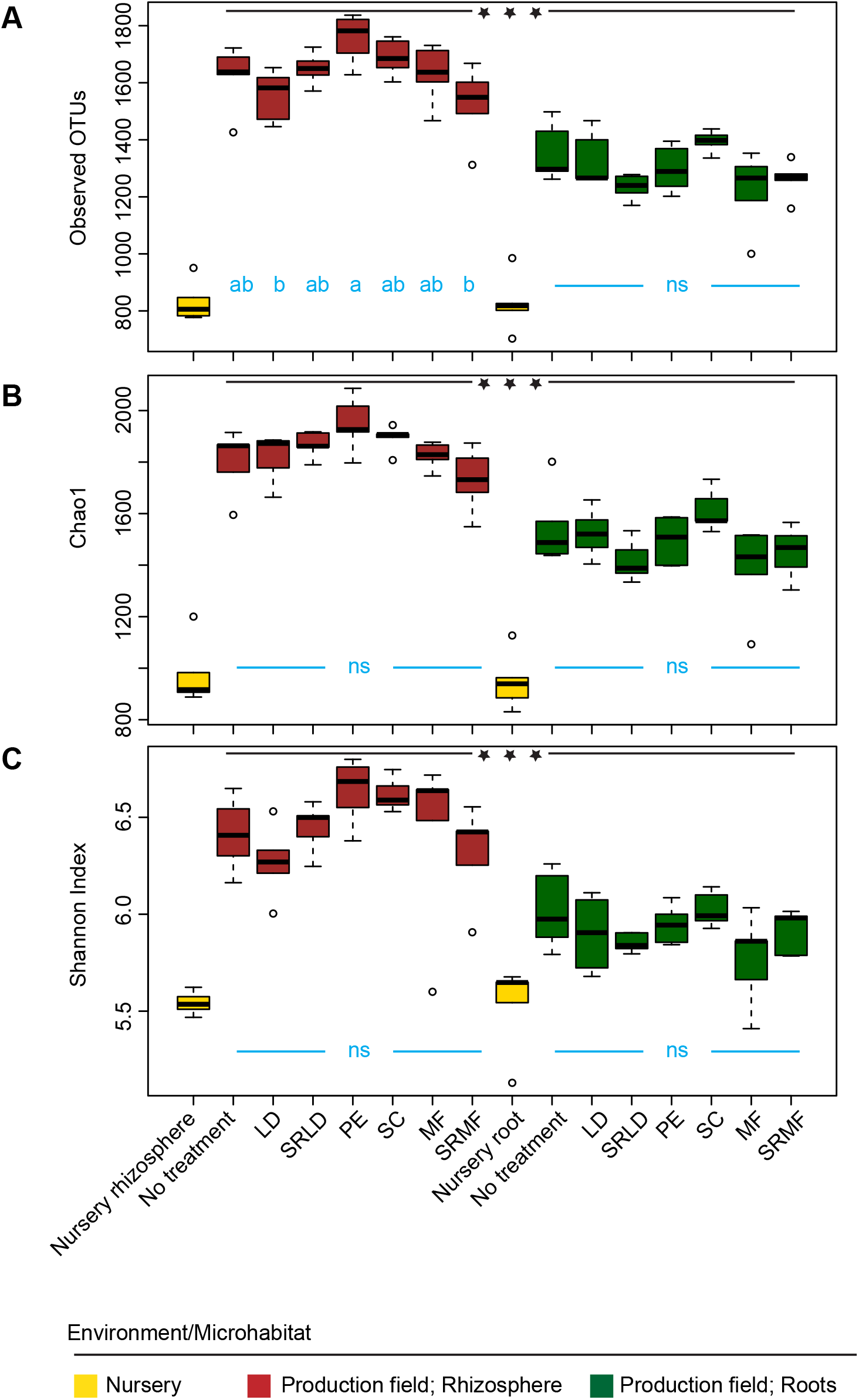
The tomato root microbiota is a gated community. Average **(A)** number of observed OTUs, **(B)** Chao 1 index and **(C)** Shannon index computed on the indicated rhizosphere and root specimens. Abbreviations LD (Liquid Digestate), SRLD (Slow acting Liquid Digestate), PE (Pelleted Digestate); SC (Organo-mineral fertiliser); MF (Mineral Fertiliser); SRMF (Slow acting Mineral Fertiliser). Asterisks denote statistically significant differences between microhabitat by non-parametric Wilcoxon rank sum test (P < 0.01). Different blue letters within individual microhabitats denote statistically significant differences between treatment means by Kruskal-Wallis non parametric analysis of variance followed by Dunn’s post-hoc test (P < 0.05); ns, no significant differences observed.

Congruently, beta-diversity analysis computed on the non-rarefied dataset using both weighted Unifrac and Bray-Curtis indicated a microhabitat-dependent microbiota diversification. In particular, the weighted Unifrac matrix visualised using a Principal Coordinates Analysis revealed such a microhabitat effect on samples processed at harvest time along the axis accounting for the major variation. Interestingly, younger nursery samples displayed a similar degree of diversification, although their communities were separated from the harvest samples on the axis accounting for the second source of variation (Figure 3). These data were supported by a PERMANOVA which attributed a R^2^ of 30% to the microhabitat, a R^2^ of 28% to the ‘Nursery/Harvest effect’ and a R^2^ of 2% to their interactions (Adonis test, 5,000 permutations, p <0.01). The analysis conducted on rhizosphere and root samples at harvest stage revealed that, congruently with the observed diversification along the axis accounting for the major variation, the microhabitat remained the major driver of the tomato communities (R^2^ 47%, Adonis test, 5,000 permutations, p <0.01) while the individual fertiliser treatments impacted these plant-associated microbial assemblages to a lesser, but significant, extent (R^2^ 13%, Adonis test, 5,000 permutations, p <0.01). This suggest that, rather than on richness per se, the fertiliser treatment impacts on the abundances and phylogenetic assignments of members of the tomato microbiota. Remarkably, the Bray-Curtis matrix produced a congruent results, although the temporal effect (i.e., nursery vs. harvest time) explained slightly more variation (~ 29%; Supplementary Figure S2) than microhabitat diversification manifested along the second axis of variation (~ 26%; Supplementary Figure S2). Crucially, also in this case the observed diversification was supported by a PERMANOVA which attributed a R^2^ of 23% to the microhabitat, a R^2^ of 29% to the ‘Nursery/Harvest effect’ and a R^2^ of 3% to their interactions (Adonis test, 5,000 permutations, p <0.01).

**Figure 3.**
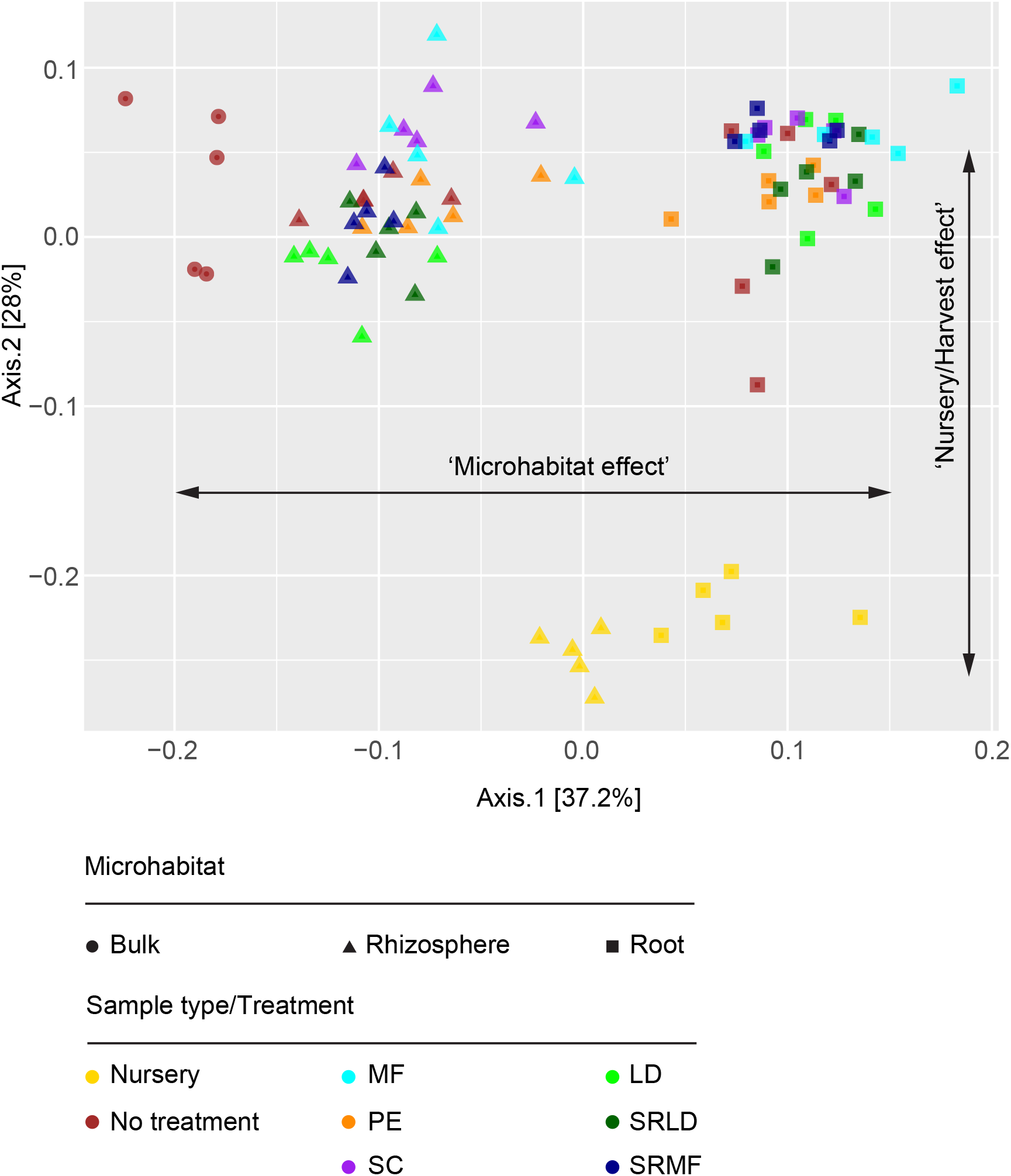
The tomato rhizosphere and root microbiota host compositionally different communities. PCoA calculated using a weighted UniFrac matrix calculated on the OTUs clustered at 97% identity among the indicated microhabitat and treatments. Abbreviations: LD (Liquid Digestate), SRLD (Slow acting Liquid Digestate), PE (Pelleted Digestate); SC (Organo-mineral fertiliser); MF (Mineral Fertiliser); SRMF (Slow acting Mineral Fertiliser).

### Differential bacterial enrichments define microhabitat and treatment “signatures” on the field grown tomato microbiota

To gain insights into individual members of the tomato microbiota responsible for the observed diversification we implemented a series of pair-wise comparisons among microhabitats and treatments at harvest stage. We took a two-pronged approach. First, we identified bacteria underpinning the microhabitat effect i.e., the selective enrichment of bacteria in the roots and the rhizosphere microhabitats amended with no fertiliser. Next, we assessed the effect of the fertiliser treatment on roots and rhizosphere bacterial composition by comparison with bacteria enriched in untreated samples.

This allowed us to identify 170 bacterial OTUs whose abundance was significantly enriched in and differentiated between rhizosphere specimens and unplanted soil samples (Wald test, p <0.01, FDR corrected; Supplementary database 1). Similarly, we identified 374 bacterial OTUs whose abundance was significantly enriched in and differentiated between root specimens and unplanted soil samples (Wald test, p <0.01, FDR corrected; Supplementary database 1). Of these differentially enriched bacteria, 96 OTUs represented a set of tomato-competent OTUs capable of colonising both the rhizosphere and root environments. When we then looked into the taxonomic affiliations of this tomato-competent microbiota, we discovered that it is dominated by members of Actinobacteria, Bacteroidetes, Alpha-, Beta-, Gamma- and Deltaproteobacteria as well as Verrucomicrobia (Figure 4). Strikingly, the taxonomic investigation revealed a bias for Actinobacteria in the root compartment, possibly reflecting an adaptive advantage of members of this phylum in colonising the endophytic environment.

**Figure 4.**
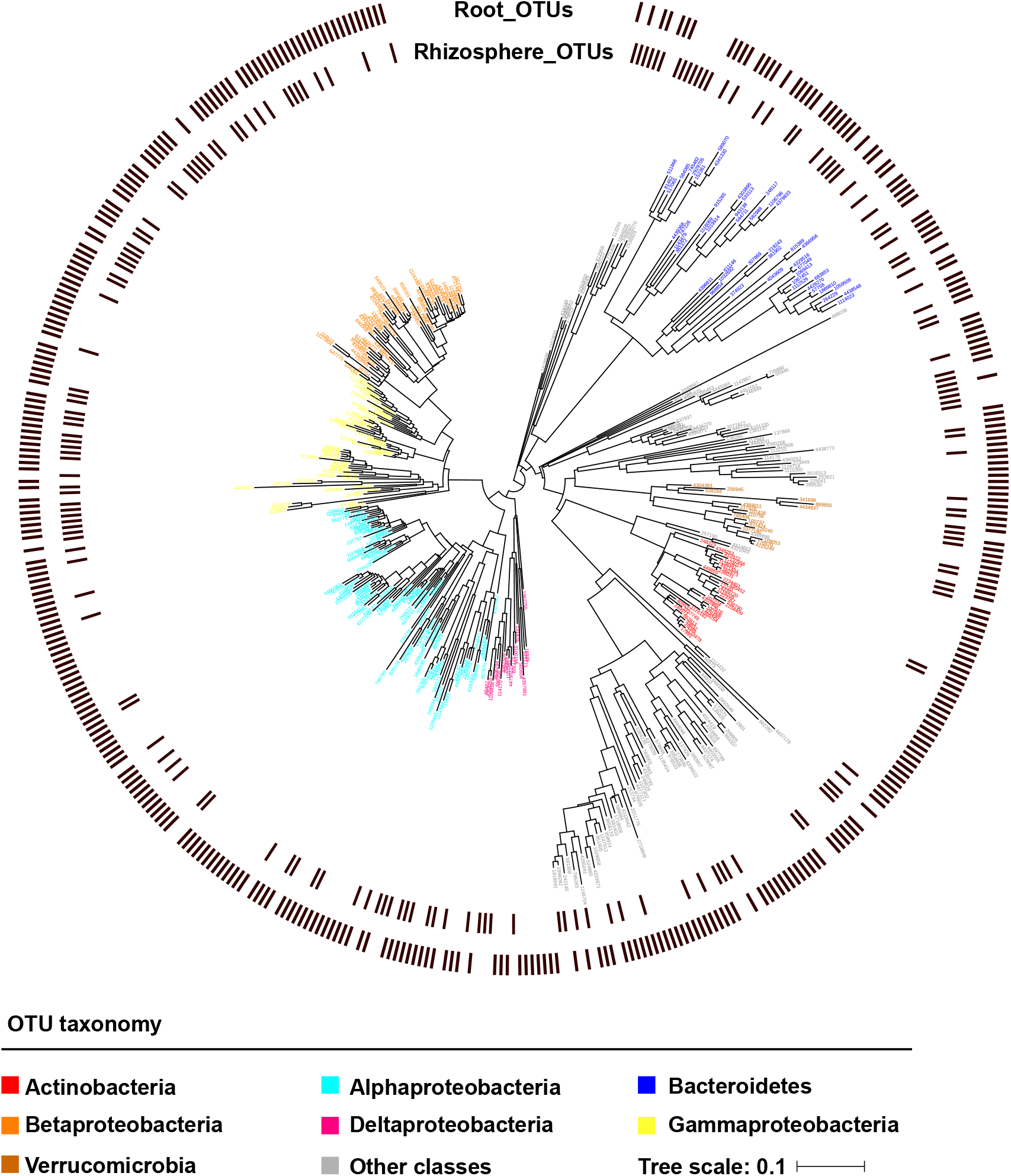
The enrichment of Actinobacteria is a distinctive feature of the tomato root microbiota. Phylogenetic relationships of the OTUs enriched in rhizosphere and root compartment. Individual external nodes represent one of the OTUs enriched in either (or both) rhizosphere or root samples in no treatment conditions (Wald test, P value < 0.01, FDR corrected) whose colour reflects their taxonomic affiliation at Phylum level. A black bar in the outer rings depicts whether that given OTU was identified in the rhizosphere- or root-enriched sub-communities, respectively. Phylogenetic tree constructed using OTUs 16S rRNA gene representative sequences.

Interestingly, each fertiliser treatment had a distinct impact on these tomato-enriched microbiota. The pelleted digestate (PE) and the slow-acting synthetic fertiliser (SRMF) yielded the highest number of uniquely enriched OTUs regardless of the microhabitat investigated, albeit with a distinct pattern: the SRMF had a more pronounced effect on the rhizosphere communities while the PE impacted more on the bacteria thriving in association with root tissues. (Wald test, p <0.01, FDR corrected; Figure 5; Supplementary database 1).

**Figure 5.**
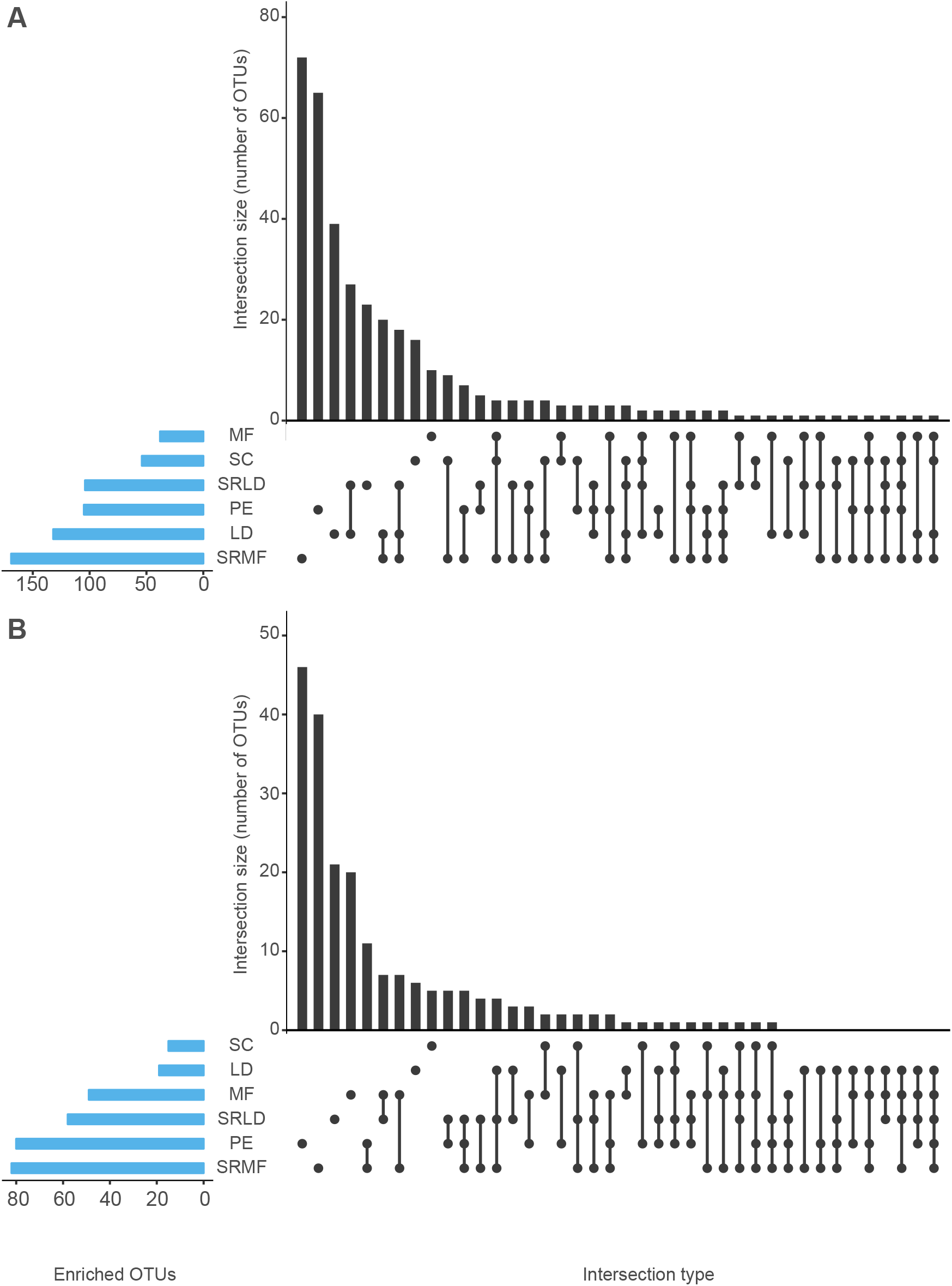
Nitrogen fertiliser modulates bacterial enrichment in the tomato rhizosphere and root compartments. Number of OTUs significantly enriched (Wald test, P value < 0.01, FDR corrected) in the indicated treatment versus untreated controls in (**A**) rhizosphere and (**B**) roots. In each panel, blue bars denote the total number of enriched OTUs for a given treatment, the black bars denote the magnitude of the enrichment in either the individual treatment or among two or more treatments highlighted by the interconnected dots underneath the panels. Abbreviations: LD (Liquid Digestate), SRLD (Slow acting Liquid Digestate), PE (Pelleted Digestate); SC (Organo-mineral fertiliser); MF (Mineral Fertiliser); SRMF (Slow acting Mineral Fertiliser).

Interestingly, when we inspected the taxonomic composition of the bacteria differentially impacted by the fertiliser treatment we observed an increase of the number of OTUs belonging to phylum of Actinobacteria. In particular, PE had 12 OTUs out of 80 and 14 OTUs out of 105, in root and rhizosphere, respectively, belonging to phylum Actinobacteria. While, MF had 15 OTUs out of 38 and 22 OTUs out of 49 in root and rhizosphere, respectively, belonging to phylum Actinobacteria (Supplementary database S1). Within this phylum we observed the presence of OTUs classified as *Streptomyces spp., Agromyces sp., Microbispora sp*. and *Actinoplanes spp*.

Together these data suggested that the enrichment of specific bacteria underpins the observed microhabitat effect whose magnitude is fine-tuned by the applied fertiliser.

### Organic- and synthetic-based fertiliser trigger different metabolic capacities in the tomato root microbiota

To investigate the ecological significance of the observed differential recruitments among fertiliser treatment we employed a predictive metagenomics approach. Briefly, we inferred *in silico* the functions encoded by the tomato microbiota at harvest stage (Materials and Methods) and we grouped the samples in digestate-based (i.e., PE, LD and SRLD; hereafter ‘organic’) and treatments containing at least a synthetic component (i.e., SC, MF and SRMF; hereafter ‘mineral’). We observed that the functions putatively encoded by the communities exposed to either organic or mineral fertilisers can discriminate between treatments in both microhabitats (PERMANOVA: Rhizosphere samples R^2^ = 14%, p value <0.01, 5,000 permutations; Root samples R^2^ = 16%, p value <0.01, 5,000 permutations). Congruently, we identified a set of 14 functions differentially enriched between root communities exposed to either group of treatments (Welch t-test, p <0.01, FDR corrected; Figure 6). Interestingly, we observed a striking dichotomy between the two groups of treatments: communities exposed to mineral fertilisers are predicted to enrich for genes implicated in the ABC transporter machinery while bacteria exposed to the organic treatments are predicted to enrich for genes implicated in the two-component system. These two set of genes are dominant in communities exposed to both treatments and are also associated to additional distinct enrichment patterns, most notably including nitrogen metabolism (organic communities) and tetracycline biosynthesis (mineral communities).

**Figure 6.**
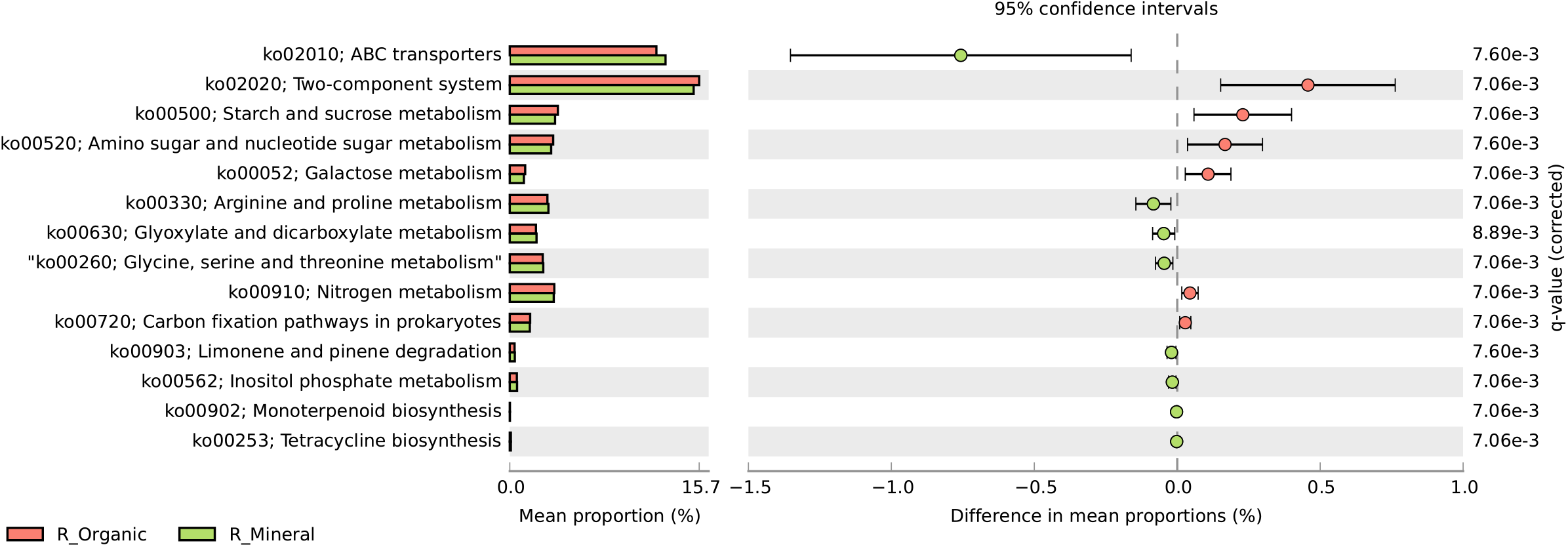
Digestate- and mineral-based fertilisers trigger a functional diversification of the tomato root microbiota. Prokaryotic functions discriminating between Digestate-based (indicated as ‘R_organic’: Liquid Digestate; Slow acting Liquid Digestate and Pelleted Digestate) and mineral-based fertilisers (indicated as ‘R_mineral’: Organo-mineral; Mineral Fertiliser and Slow acting Mineral Fertiliser) retrieved from Tax4Fun functional profiles (Welch’s t-test FDR corrected, p<0.01).

These results suggest that, within tomato roots, the observed taxonomic diversification underpins a functional specialisation of the microbiota which, in turn, may impact on plant growth development and health.

## DISCUSSION

This study revealed that all nitrogen treatments led to an increase of tomato production in comparison with the no fertilization treatment (fold change between 0.8 and 1.73) confirming that, in the tested conditions, nitrogen limits the yield potential of processing tomato crops as observed in previous studies (Ronga et al., 2015; Ronga et al., 2017). Yet, despite the same amount of nitrogen was applied in each treatment (i.e., 150 kg ha^−1^), all the treatments were statistically different from each other. A prediction of this observation is that, under the tested conditions, the nature of the fertilisers, rather than the amount of nitrogen per se, affect the yield and the fruit quality of tomato plants. These observations and the putative contribution to fertiliser use efficiency of the microbial communities thriving at the root-soil interface (Alegria Terrazas et al., 2016), motivated us to investigate relationships between yield traits and the composition of the tomato rhizosphere and root microbiota under field conditions.

### The tomato rhizosphere and root microbiota are gated communities

First, we characterised the rhizosphere and root microbiota of processing tomato with no treatment. Both alpha and beta diversity discriminated between the communities of seedlings and adult plants. Despite these differences, which could be attributed to both abiotic, e.g., time of residence in soil (Dombrowski et al., 2017), and biotic factors, e.g., developmental-conditioned rhizodeposits (Chaparro et al., 2014), it is striking to note how tomato plants displayed a rhizosphere and root compartmentalisation regardless of the developmental stage. This is congruent with the observation that in rice, the assembly and structural diversification of the microbiota is a rapid process which reaches a steady-state level within a few weeks from germination (Edwards et al., 2015). Closer inspection of the rhizosphere and root profiles at harvest stage indicates that these plant-associated communities are phylogenetically related to those of unplanted soil, suggesting that the initial tomato microbiota is further modulated by the growing conditions.

Despite this apparent relatedness, the selective enrichment of individual bacterial members of the microbiota discriminates between rhizosphere and root communities for mature plants from unplanted soil profiles (Figure 4). These enrichments displayed a bias for members of the phyla Actinobacteria, Bacteroidetes, Proteobacteria (including the classes Alpha--, Beta-, Delta- and Gammaproteobacteria) as well as Verrucomicrobia. Members of these taxa have routinely been reported in studies focussing on plant-competent bacteria under both laboratory and field conditions (Bulgarelli et al., 2013; Walters et al., 2018), suggesting that the experimental approach followed in this study can be considered representative for field-grown processing tomato.

However, we noticed a differential selective pressure on the bacteria thriving either in the rhizosphere or in the root tissue: this latter environment produced more distinct profiles, i.e. more differentially enriched bacteria compared to unplanted soil, than the ones retrieved from the soil surrounding the roots. This indicates that the diversification of the tomato-inhabiting microbial communities from the surrounding soil biota initiates in the rhizosphere and progresses through the root tissue, where it produces a more pronounced microbiota diversification compared to unplanted specimens. This observation is reminiscent of the recruitment patterns of other crops such as barley (Bulgarelli et al., 2015) but it is in striking contrast with studies conducted with both model (Bulgarelli et al., 2012) and field-grown (Rathore et al., 2017) Brassicaceae, whose ‘rhizosphere effect’ appears negligible.

We further noticed that the “root effect” on the microbiota was exerted also at phylogenetic level with a bias for the enrichment Actinobacteria. This observation is in apparent contrast with results gathered from the recent seed-to-seed characterisation of the tomato microbiota which revealed that, albeit averaging 8% of the sequencing reads across microhabitats, members of this phylum did not significantly discriminate root from rhizosphere specimens (Bergna et al., 2018). However, it is worth mentioning that these two studies differed in terms of both soil type and plant genotype used.

Together, our results suggest that both species- and soil-specific traits govern the assembly of the tomato microbiota in field-grown crops.

### Nitrogen source impacts on the structural and functional composition of the tomato microbiota

Next, we investigated the impact of the type of nitrogen fertiliser on the tomato microbiota and we demonstrated that each treatment produced “distinct signatures”, represented by specific selective enrichment, on both the rhizosphere and root communities. Despite microhabitat-associated variation, the effect of the application of pelleted digestate (PE) resulted in the most distinct microbial profile in the root compartment and the second largest number of specifically enriched OTUs in the rhizosphere Of note, the slow-acting mineral fertiliser (SRMF) follow a “complementary” pattern: its application yielded the greatest and the second greatest number of differentially enriched OTUs compared to untreated samples in the rhizosphere and root profiles, respectively. Remarkably, these two treatments had a discernible effect also on crop yield, with the PE treatment producing the best performance among the various fertilisers. Our data are congruent with studies conducted on wheat which observed a structural diversification of the soil and plant-associated communities exposed to either mineral or organic fertilisers (Kavamura et al., 2018). Yet, the numerical shift in terms of OTUs differentially enriched per se cannot explain the potential impact of these communities on crop yield: owing to the fact that the SMRF treatment, which is associated to a significant reduction in yield traits (compared to PE) is capable of triggering a comparable OTU enrichment.

We therefore focused our attention on the taxonomical composition of the rhizosphere and root communities. In particular, we noticed that the proliferation of Actinobacteria in the root compartment was retained in the various treatments. The enriched Actinobacteria included *Streptomyces spp., Agromyces sp., Microbispora sp*. and *Actinoplanes spp*. (Supplementary Database 1). *Streptomyces spp*. are well-known bacteria able to produce a wide diversity of bioactive compounds able to promote plant growth and health (de Jesus Sousa and Olivares, 2016). On the other hand, members of the genus Streptomyces are responsible of economically relevant plant diseases, most notably common scab of potato caused by *S. scabies* (Loria et al., 2006).

Thus, the taxonomic diversification triggered by both microhabitat and treatment may underpin a functional diversification of the microbiota at the cross-road of mutualism and inter-species competition.

This functional diversification of the root communities is manifested by the differential enrichments of ABC transporter genes (mineral) and the two-component system (organic). Although predictive metagenomics is inherently limited by fact that the individual phylogenetic marker used (i.e., the 16S rRNA gene) may fail to recapitulate the genetic diversity existing among strains of the same phylogenetic lineage (Karasov et al., 2018), ABC transporters have previously been identified as genes underpinning rhizosphere competence in the microbiota of wheat and cucumber (Ofek-Lalzar et al., 2014). Likewise, the two-component system is required for the rhizosphere colonisation of the biocontrol agent *Pseudomonas fluorescens* WCS365 (De Weert et al., 2006). These observations indirectly support the results gathered from our predictive metagenomics approach. Owing the role played by these classes of genes in uptake of organic compounds (e.g., root exudates, cellular secretion) and stimulus-response mechanisms (e.g., chemotaxis) respectively, it is tempting to hypothesize that the different source of nitrogen define a different metabolic status in and in the vicinity of tomato roots which, in turn, requires a prompt adaptation of the root-inhabiting communities.

For instance, experimental data indicate that the abundance of phytoavailable nitrogen, i.e., the scenario of mineral fertiliser treatments, tends to repress the proliferation and activity of members of the microbiota (Ramirez et al., 2012; Terrazas et al., 2019), and this in turn may be reflected in the metabolism of secondary compounds (terpenoid and polyketide metabolism) and membrane transport (ABC transporters).

A “true” comparative metagenomics investigation, whereby the individual communities are subjected to shot-gun sequencing, will be ultimately necessary to test these hypotheses.

We further hypothesize that this adaptation is modulated by mineral nitrogen availability, as manifested by the differential enrichment of functions associated to nitrogen metabolism *per se* and aminoacids. This observation is congruent with results gathered from monocots wheat (Kavamura et al., 2018) and rice (Zhang et al., 2019) and suggests a cross-species pattern whereby plant’s adaptation to nitrogen forms and availability is mediated, at least in part, by the associated microbiota.

Finally, it is interesting to note how the production of antibiotics, namely tetracyline, is also among the functions differentially enriched between fertilisers. It is becoming increasingly clear how plant-associated bacteria can act as a reservoir of antimicrobial genes (Cernava et al., 2019) which can be deployed during inter-organismal competition in the plant microbiota. This hypothesis could be tested by leveraging on indexed- and genome-annotated bacterial collection for the tomato microbiota, similar to the approach pursued with bacteria isolated from other plant species (Levy et al., 2018).

Our investigation suggests that the bacterial microbiota of field-grown processing tomato is the product of a selective process that progressively differentiates between rhizosphere and root microhabitats. This process initiates as early as plants are in a nursery stage and it is then more marked when plants reached the harvest stage. This selection *a*) acts both on the relative abundances and phylogenetic assignments of members of the tomato microbiota, *b*) is modulated, at least in part, by the nitrogen fertiliser provided which, in turn, *c*) triggers different microbial metabolic specialisations within tomato roots.

It is important to mention that the nitrogen fertiliser may also represent a microbial inoculant *per se*, in particular in the case of organic-based amendments. For instance, a comparative study of 29 different full-scale anaerobic digestion installations revealed that Firmicutes, followed by Bacteroidetes and Proteobacteria, dominated the resulting microbial communities (De Vrieze et al., 2015). Considering the plant-associated profiles observed in this study, in particular the enrichment of Actinobacteria in the root communities, it is legitimate to hypothesize that the input digestate bacteria may act as in inoculum for a part of the tomato microbiota, which is further fine-tuned by the exposure to soil microbes. Future studies, integrating the microbial profiling of the input fertiliser treatment, will be required to accurately elucidate microbial dynamics associated with the application synthetic (i.e., germ-free) and organic fertilisers.

### Towards a lab-in-the-field approach to harness the potential of plant microbiota for climate-smart agriculture

Our experiments represent an example of how cultivation-independent approaches can be efficiently deployed to investigate the plant microbiota under field conditions. Although this type of investigation is not novel *per se* in tomato (Toju et al., 2019), our results revealed fundamentally novel insights into plant’s adaptation to nitrogen fertilisers and the implication for crop yield. Similar to what has recently been postulated for tomato pathogen protection (Kwak et al., 2018), our results predicts that the use of field-derived, sequencing data will allow scientists to identify “signatures” of the plant microbiota that can be targeted to enhance plant performance. This approach, which we define as lab-in-the-field, will be key towards the rationalisation of nitrogen (and other treatments) application in agriculture and we anticipate will pave the way for the effective exploitation of the plant microbiota for agricultural purposes (Schlaeppi and Bulgarelli, 2015; Toju et al., 2018).

## Supporting information

Supplementary Database 1

## AUTHORS CONTRIBUTION

DR conceived of and designed the field experiment. DR and FC harvested the field data and samples. FC, RAT, EF and DB conceived of and designed the analysis of the microbiota. FC, MC, CVGA, SR-A performed the microbiota experiments. FC, RAT and DB analysed the sequencing data. FC, RAT and DB wrote the initial draft of the manuscript. All the authors discussed the results and commented on the manuscript.

## ACKNOWLEDGEMENTS

We thank Dr Guido Bezzi (CIB - Italian Biogas Association, Italy) for providing us with the field, plant material and the digestate treatments, Dr. Massimo Zaghi (CAT - COOP. Agroenergetica Territoriale Correggio S.C.A., Italy) for providing the pelleted digestate and Dr Stefano Tagliavini (SCAM Spa, Italy) for providing the organo-mineral fertiliser.

We thank Malcolm Macaulay, Jenny Morris and Dr Pete Hedley (The James Hutton Institute, Invergowrie, UK) for their support in preparing the sequencing library and generating the sequencing data.

We thank Prof Marco Candela (University of Bologna, Italy) for the critical comments on the manuscript.

