## Supplementary figures and images for "Nitrogen Fertilisers Shape the Composition and Predicted Functions of the Microbiota of Field-Grown Tomato Plants"

### Supplementary Figure S1

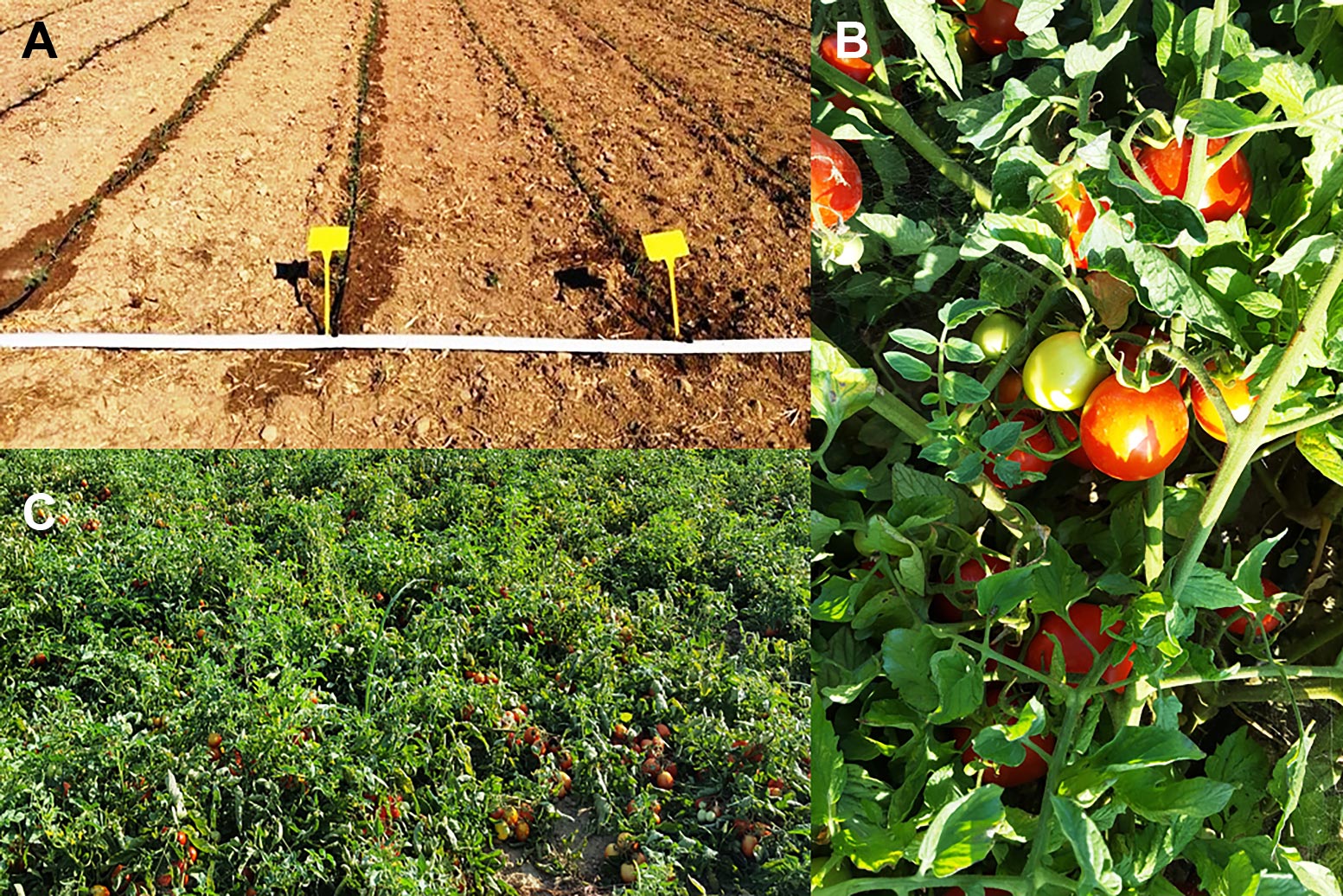

### Supplementary Figure S2

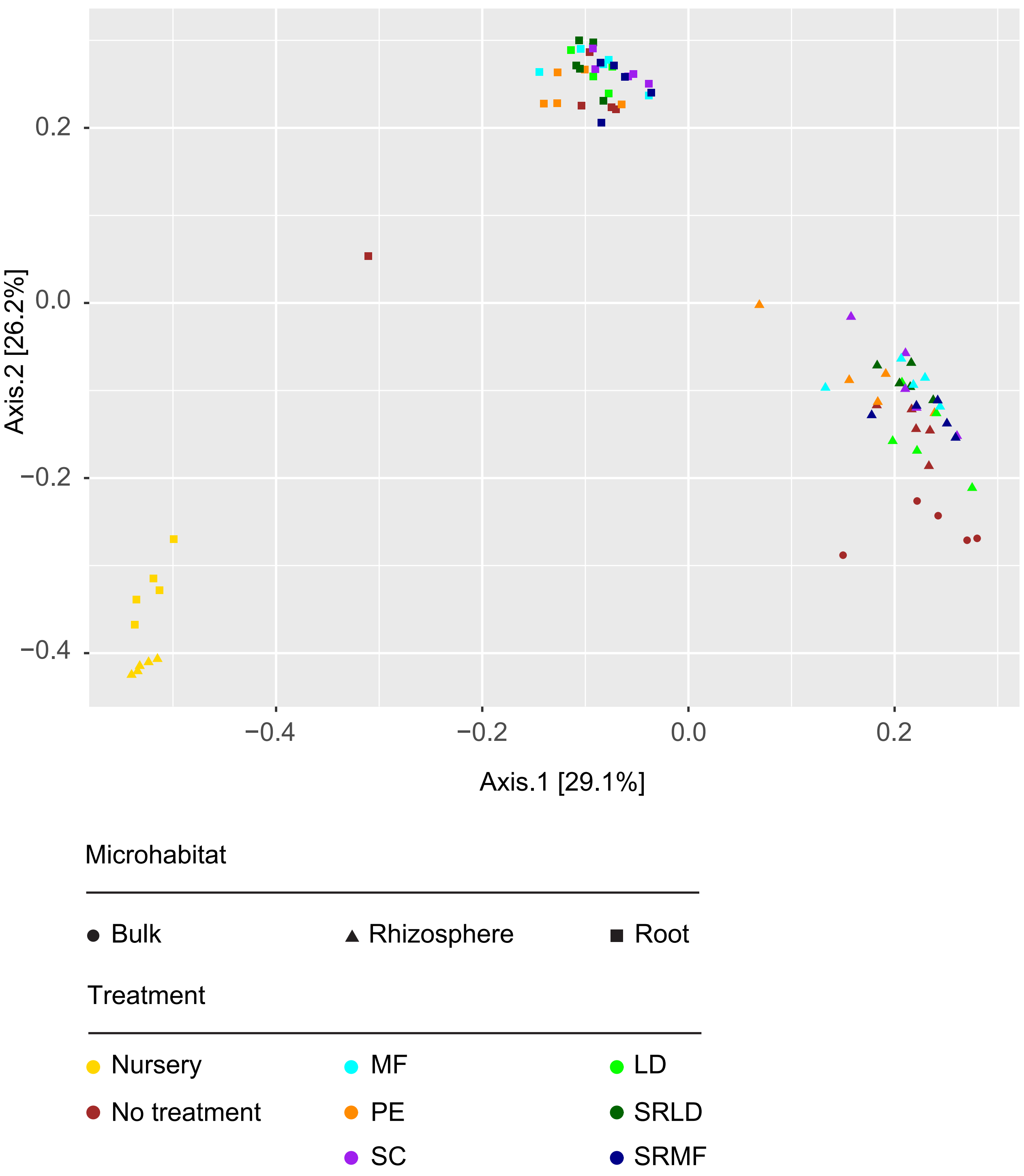
